# Boosting pro-vitamin A content and bioaccessibility in leaves by combining engineered biosynthesis and storage pathways with high-light treatments

**DOI:** 10.1101/2023.05.22.541812

**Authors:** Luca Morelli, Pablo Perez-Colao, Diego Reig-Lopez, Xueni Di, Briardo Llorente, Manuel Rodriguez-Concepcion

**Affiliations:** Institute for Plant Molecular and Cell Biology (IBMCP), CSIC-Universitat Politècnica de València, 46022 Valencia, Spain; ARC Center of Excellence in Synthetic Biology, Australian Genome Foundry, and School of Natural Sciences, Macquarie University, Sydney, NSW, 2109, Australia

**Keywords:** Carotenoids, bioaccessibility, biofortification, β-carotene, lettuce, plastoglobules, vitamin A

## Abstract

The relevance of plants as food is expected to grow for a more sustainable diet. In this new context, improving the nutritional quality of plant-derived foods is a must. Biofortification of green leafy vegetables with pro-vitamin A carotenoids such as β-carotene has remained challenging to date. Here we combined two strategies to achieve this goal. One of them (that we call strategy C) involves producing β-carotene in extraplastidial locations of leaf cells to avoid the negative impacts on photosynthesis derived from changing the balance of carotenoids and chlorophylls in chloroplasts. The second approach (that we refer to as strategy P) involves the conversion of chloroplasts into non-photosynthetic, carotenoid-overaccumulating chromoplasts in some leaves, leaving other non-engineered leaves to sustain normal plant growth. Combination of these two strategies resulted in a 5- fold increase in the amount of β-carotene in *Nicotiana benthamiana* leaves. Following several attempts to further improve β-carotene leaf contents by metabolic engineering, hormone treatments and genetic screenings, it was found that promoting the proliferation of plastoglobules with high-light treatments not only improved β-carotene accumulation but it also resulted in a much higher bioaccessibility. Combination of strategies C and P together with a high-light treatment increased the levels of accessible β-carotene 30-fold compared to controls. We further demonstrate that stimulating plastoglobule proliferation with strategy P but also with high-light alone can also stimulate and hence improve β-carotene contents and bioaccessibility in edible lettuce (*Lactuca sativa*) leaves, unveiling the power of non-GMO approaches for leaf biofortification.

## INTRODUCTION

Micronutrient deficiency, also known as hidden hunger, is still a major problem in many countries. In particular, vitamin A deficiency causes xerophthalmia and can lead to other health problems and even death, affecting children from malnourished populations worldwide. The incorporation of micronutrients as dietary supplements or as food ingredients (i.e., food supplementation or fortification, respectively) can be a solution, but these strategies remain unaffordable in many cases (Fitzpatrick et al., 2012). An alternative approach is the development of crop varieties enriched in micronutrients, i.e., biofortification (Morelli and Rodriguez-Concepcion, 2023). Among biofortification strategies, biotechnology has the highest potential for a fast and focused enrichment of the desired plant tissues. In the case of vitamin A, biofortification of rice with β-carotene (the main pro-vitamin A carotenoid) was implemented to create the widely known Golden Rice (Ye et al., 2000; Paine et al., 2005). After continuous development to further increase β-carotene contents and to include this trait into local rice varieties, this is widely recognized as an example of successful biofortification despite the controversies arising from the use of GMOs (Welsch and Li, 2022).

Most successful strategies for carotenoid biofortification have been reported in non-photosynthetic tissues (e.g., canola seeds, potato tubers, or rice endosperm). In fact, manipulation of carotenoid levels in photosynthetic tissues is much more challenging. In photosynthetic systems, carotenoids participate in light harvesting and are essential for photoprotection, whereas they play a non-essential role as pigments ranging from yellow to red in non-photosynthetic tissues and organisms (Lichtenthaler, 2012; Rodriguez-Concepcion et al., 2018). Chloroplast carotenoid profile is quite similar in most plant species - with β-carotene accounting for about 20% of total carotenoids - and both composition and levels must be finely balanced with those of chlorophylls for an efficient assembly and functionality of photosynthetic complexes (Domonkos et al., 2013; Esteban et al., 2015; Hashimoto et al., 2018). Low levels of carotenoids are present in the chloroplast envelope membrane, while most carotenoids are associated with proteins in various photosynthetic complexes located in the thylakoid membranes. Specifically, β-carotene is primarily associated with photosystems (PSI and PSII) and the cytochrome b6f complex. An increase in β-carotene levels in chloroplasts may hence interfere with plant photosynthesis (Pascal et al., 2005; Lichtenthaler, 2012; Domonkos et al., 2013).

Unlike chloroplasts, the carotenoid-accumulating plastids found in pigmented non-photosynthetic tissues - called chromoplasts – have a very diverse carotenoid composition. Chromoplasts naturally differentiate from different types of plastids, including leucoplasts, amyloplasts, or, most frequently, chloroplasts (Sun et al., 2018; Sadali et al., 2019). Recent results have demonstrated that leaf chloroplasts can be artificially converted into chromoplasts by overexpressing a bacterial phytoene synthase enzyme - crtB - that catalyzes the first committed step of the carotenoid pathway (Llorente et al., 2020; Torres-Montilla and Rodriguez-Concepcion, 2021). Artificial leaf chromoplasts accumulate higher levels of carotenoids and are particularly enriched in β-carotene (Llorente et al., 2020; Morelli et al., 2023). The formation of chromoplasts (natural or artificial) involves the development of internal structures that sequester and store carotenoids. These structures differ among chromoplasts depending on the carotenoid content, tissue, or plant species. For example, tomato fruit chromoplasts and crtB-generated leaf chloroplasts accumulate β-carotene in plastoglobules (PG), whereas carrot root chromoplasts accumulate much higher concentrations of β-carotene in crystalline bodies (Morelli et al., 2023; Kim et al., 2010; Schweiggert et al., 2012).

In plant cells, carotenoids are synthesized from the five-carbon (C_5_) isoprenoid precursors isopentenyl diphosphate (IPP) and its isomer dimethylallyl diphosphate (DMAPP) produced by the plastidial methylerythritol 4-phosphate (MEP) pathway (Rodríguez-Concepción and Boronat, 2002). Condensation of three IPP and one DMAPP molecules forms C_20_ geranylgeranyl diphosphate (GGPP), the direct precursor for carotenoids and other plastidial isoprenoids such as diterpenes, chlorophylls, tocopherols, plastoquinones and phylloquinones (Figure 1). The first committed step of the carotenoid pathway is the formation of C_40_ phytoene from two molecules of GGPP. This reaction is catalyzed by the enzyme phytoene synthase (PSY in plants, crtB in bacteria). Colorless phytoene is then sequentially desaturated and isomerized to lycopene, the red pigment typical of ripe tomato fruit. Cyclization of the two ends of the linear lycopene molecule generates β-carotene (with two β rings) or α-carotene (with one ε ring and one β ring). The formation of two ε rings is very rare in plant carotenoids, although there are examples in crops such as lactucaxanthin in lettuce (Britton, 1995; Phillip and Young, 1995). Oxidative modification of the rings generates oxygenated carotenoids - known as xanthophylls - such as violaxanthin and neoxanthin (from β-carotene) or lutein (from α-carotene, an intermediate that is hardly detected in chloroplasts). The same precursors used for carotenoid biosynthesis in plastids, IPP and DMAPP, are also produced in the cytosol by the mevalonate (MVA) pathway (Rodríguez-Concepción and Boronat, 2015). MVA-derived IPP and DMAPP are mostly used to produce polyterpenes, sesquiterpenes, and triterpenes such as phytosterols (the major product of the pathway), while very low levels of GGPP are synthesized for the biosynthesis of diterpenes and protein prenylation. Interestingly, cytosolic IPP and DMAPP can be used to produce carotenoids by introducing bacterial or fungal enzymes that produce GGPP and convert downstream carotenoids (Figure 1) (Majer et al., 2017; Andersen et al., 2021; Zheng et al., 2023). This strategy preserves the chloroplast carotenoid profile without disrupting the photosynthetic complexes and allows the accumulation of carotenoid intermediates in the cytosol by separating them from the endogenous (plastidial) enzymes that convert them into downstream carotenoid end-products. Here we explored several strategies to biofortify leaves with β-carotene by combining its extraplastidial production and its enrichment in the PG of artificial chromoplasts.

**Figure 1.**
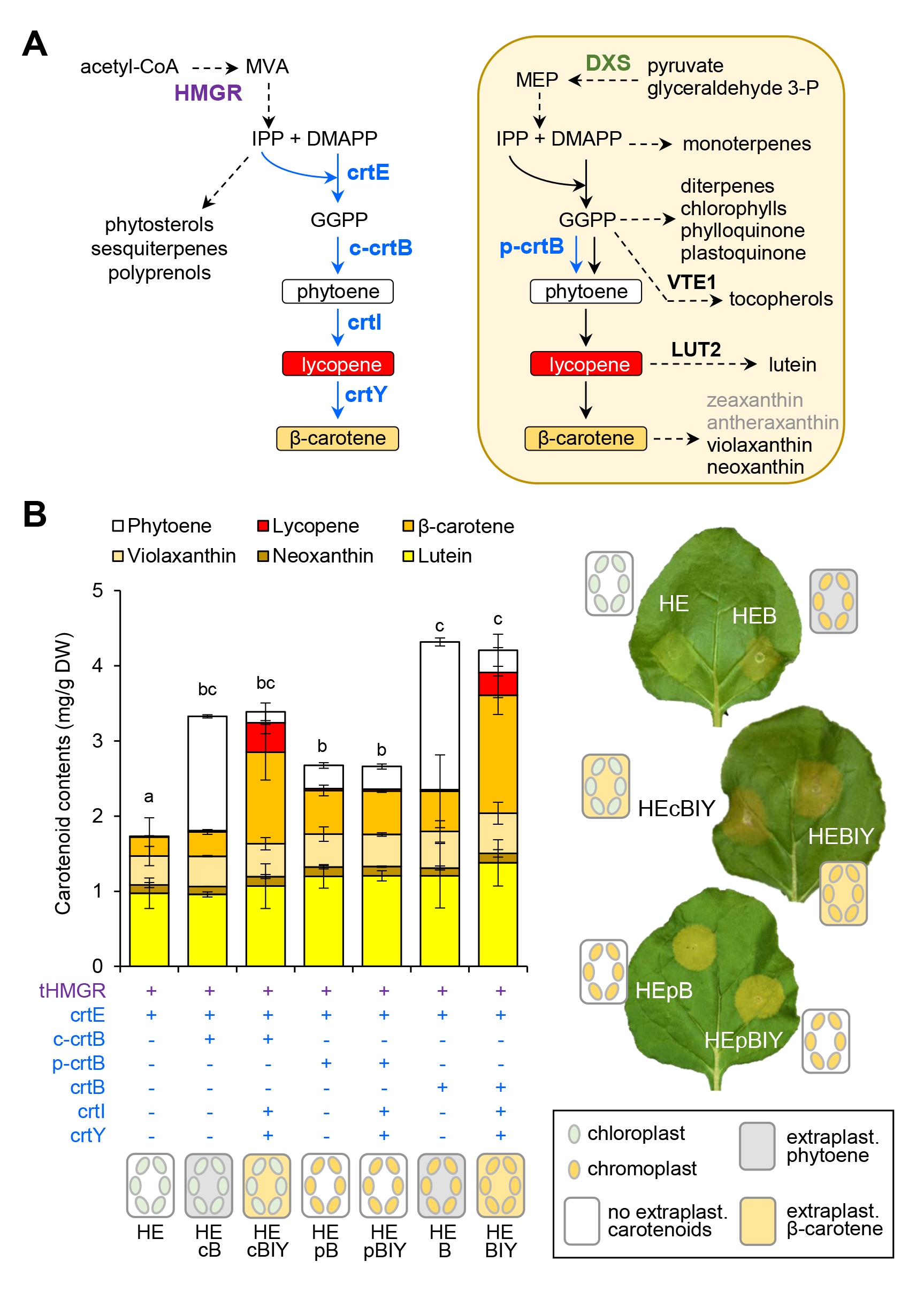
Strategies to produce β-carotene in different cell compartments. **(A)** Schematic representation of the carotenoid biosynthetic pathways and enzymes covered in this work. Dashed arrows represent multiple steps. The plastidial pathway is boxed in orange. **(B)** Carotenoid levels in *N. benthamiana* leaves agroinfiltrated with the indicated combinations and collected at 6 dpi. Mean and SD of n≥3 independent biological replicates are shown. DW, dry weight. Letters represent statistically significant differences (*P* < 0.05) among means of total carotenoid levels according to post hoc Tukey’s tests run after one way ANOVA detected different means. Representative images of leaves agroinfiltrated with the indicated combinations are also shown. Subcellular sites of carotenoid accumulation are indicated in the cartoons.

## RESULTS

### An engineered cytosolic pathway for β-carotene production

Despite the relatively high amount of β-carotene in green (leaf) tissues, its physical and functional association with the photosynthetic machinery prevents its overaccumulation. To enhance β-carotene contents in leaves without directly interfering with photosynthesis, we used an engineered cytosolic pathway previously shown to produce very high levels of extraplastidial lycopene (Andersen et al., 2021). Conversion of lycopene into β-carotene was accomplished by adding the *Pantoea ananatis crtY* gene encoding the bacterial lycopene-β-cyclase (Figure 1A). *Nicotiana benthamiana* leaves were agroinfiltrated with the resulting construct, harboring the *P. ananatis* genes encoding crtE (to produce GGPP), crtB (to transform GGPP into phytoene), crtI (to synthesize lycopene from phytoene) and crtY (to convert lycopene into β-carotene). The crtB protein in its natural conformation can localize to both cytosol and plastids (Andersen et al., 2021; Llorente et al., 2020). Therefore, a version of the enzyme with GFP fused to its N-terminus (referred to as GFP-crtB or c-crtB) was used to retain the protein in the cytosol of agroinfiltrated leaf cells (Andersen et al., 2021; Llorente et al., 2020). Furthermore, a truncated version of hydroxymethylglutaryl CoA reductase (tHMGR) was used to increase the MVA pathway flux and produce extra IPP and DMAPP for GGPP synthesis (Figure 1B) (Andersen et al., 2021; Zheng et al., 2023). Following agroinfiltration with this gene combination (named HEcBIY), samples were collected at 6 days post-infiltration (dpi) for HPLC analysis of carotenoid levels (Figure 1B). Parallel experiments were carried out with combinations of tHMGR + crtE (HE) and tHMGR + crtE + c-crtB (HEcB). Total carotenoid levels were more than doubled when c-crtB was included in the combination (Figure 1B). As expected, the main carotenoids were phytoene and β-carotene in HEcB and HEcBIY leaves, respectively (Figure 1B). The accumulation of β-carotene in HEcBIY leaves (the sum of chloroplast levels plus engineered cytosolic contents) was about 4-fold higher than the levels measured in HE leaves (corresponding to only chloroplast levels) (Figure 1B). The relatively high levels of phytoene and lycopene detected in these samples, however, indicate an incomplete conversion. The relatively high amount of lycopene remaining in HEcBIY leaf sectors explains their reddish color (Figure 1B).

### Cytosolic production of carotenoids can be combined with chromoplast development for further carotenoid enrichment of leaves

After successfully demonstrating that it is possible to produce and accumulate a large amount of β- carotene in extraplastidial locations, we tested whether this strategy (that we called strategy C) could be combined with the crtB-mediated differentiation of leaf chloroplasts into β-carotene-enriched chromoplasts (Llorente et al., 2020, Morelli et al., 2023). Artificial chromoplastogenesis (that we name here strategy P) requires crtB acting in plastids. Indeed, agroinfiltration of *N. benthamiana* leaves with combinations containing a plastid-targeted crtB enzyme (herein referred to as p-crtB) led to a characteristic yellow phenotype due to an increased accumulation of plastidial carotenoids (Figure 1B). In particular, β-carotene levels increased about 2-fold compared to control leaves lacking p-crtB. Conversion of leaf chloroplasts into chromoplasts can also be triggered by the unmodified crtB protein, which enters the plastid likely due to a cryptic transit peptide (Andersen et al., 2021; Llorente et al., 2020). Therefore, strategies C and P might be combined just by using the unmodified crtB enzyme acting in both cytosolic and plastidic compartments. A comparison of unmodified crtB, c-crtB, or p-crtB in combination with tHMGR and crtE showed that the content of phytoene measured in leaf sectors producing the untargeted protein (HEB) was roughly a sum of the amounts obtained with the cytosolic (HEcB) plus the plastidial (HEpB) enzymes (Figure 1B). An additive phenotype was also observed for β- carotene when comparing HEBIY with HEcBIY plus HEpBYI (Figure 1B). The levels of β-carotene in HEBYI samples increased about 5-fold compared to HE controls (Figure 1B).

### Providing more plastidial precursors does not improve carotenoid production due to a bottleneck at the phytoene desaturation level

The addition of tHMGR to increase the MVA pathway flux was key for the success of strategy C as it allowed to boost the production of extraplastidial carotenoids (Andersen et al. 2021; Zheng et al., 2023). We therefore reasoned that increasing the metabolic flux through the MEP pathway might also contribute to improving the contents of plastidial carotenoids produced with strategy P (Figure 1A). The enzyme deoxyxylulose 5-phoshate synthase (DXS) catalyzes the first and main rate-determining step of the MEP pathway (Rodríguez-Concepción and Boronat, 2015). Three classes of DXS proteins have been reported: class 1 (DXS1) proteins typically function as house-keeping enzymes, class 2 (DXS2) are specialized isoforms that normally increase the MEP pathway flux when required for peak demands, and class 3 (DXS3) proteins lack DXS activity (de Luna-Valdez et al., 2021). To test whether increasing the levels of true DXS enzymes might positively influence the production of carotenoids by strategy P, we agroinfiltrated *N. benthamiana* leaves with constructs for crtB and different DXS sequences. Specifically, we used class 1 DXS enzymes from Arabidopsis (AtDXS, also referred to as CLA1) and tomato (SlDXS1) and a class 2 DXS from tomato (SlDXS2). AtDXS1 is the only enzyme with DXS activity in Arabidopsis (Phillips et al., 2008; Carretero-Paulet et al., 2013). Tomato SlDXS1 is the main isoform participating in carotenoid biosynthesis in tomato (also during fruit ripening, when carotenoid accumulation is boosted as chloroplasts differentiate into chromoplasts), while SlDXS2 shows lower levels of expression in most tissues but it is induced in specialized structures such as leaf trichomes in response to environmental stimuli to produce dedicated isoprenoids (Lois et al., 2000; Paetzold et al., 2010; Brand and Tissier, 2022). Of the three isoforms, only SlDXS1 led to a statistically significant increase in total carotenoid levels compared to crtB samples (Figure 2A). Such increase was mainly due to a massive accumulation of phytoene (Figure 2A), suggesting that SlDXS1 can efficiently improve the supply of carotenoid precursors that are converted into phytoene by plastid-localized crtB. Further metabolism of phytoene into downstream carotenoids by endogenous enzymes, however, appears to be compromised. Interestingly, the loss of photosynthetic activity associated with chromoplast development in crtB samples (Llorente et al., 2020) was faster in the presence of SlDXS1 but not with AtDXS1 or SlDXS2, as monitored by quantifying the effective quantum yield of photosystem II (ɸPSII) for four days following agroinfiltration (Figure 2B). This result suggests that activated phytoene overaccumulation accelerates the conversion of chloroplasts into artificial chromoplasts, supporting our previous conclusion that phytoene overproduction is the main factor priming chloroplasts to be converted into chromoplasts as downstream carotenoids are formed (Llorente et al., 2020). In agreement with the deduced existence of a limiting step for phytoene conversion into downstream carotenoids in agroinfiltrated leaf cells, no improvement in the contents of β-carotene or any other plastidial carotenoid besides phytoene was observed when SlDXS1 was incorporated to the HEBIY combination (Figure 2C).

**Figure 2.**
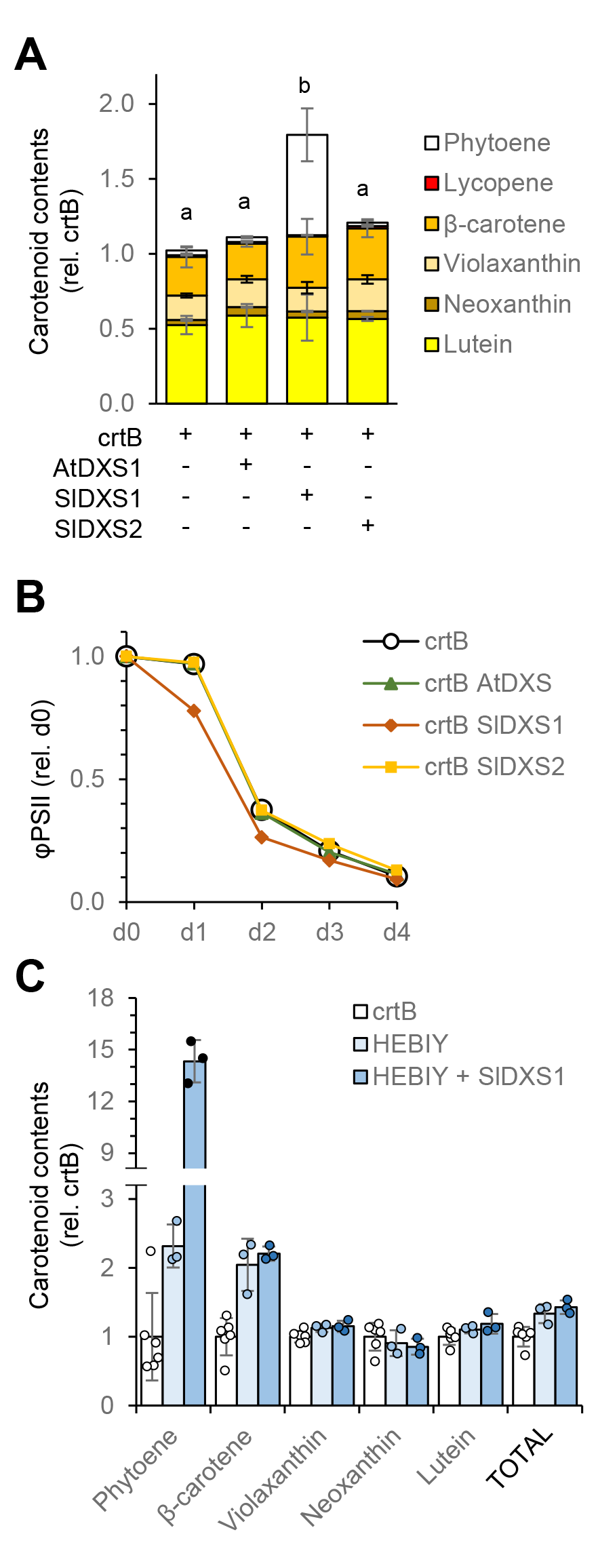
Increased supply of plastidial precursors results in phytoene overaccumulation. *N. benthamiana* leaves were agroinfiltrated with the indicated constructs and analyzed at 6 dpi. **(A)** Relative carotenoid levels shown as the mean and SD of n≥3 independent biological replicates. Letters represent statistically significant differences (*P* < 0.05) among means of total carotenoid levels according to post hoc Tukey’s tests run after one way ANOVA detected different means. **(B)** Relative effective quantum yield of photosystem II (φPSII). Mean and SD of at least 5 measurements in n≥3 independent leaf samples are expressed relative to day 0 values. **(C)** Relative levels of the indicated carotenoids represented as individual values (dots) and the corresponding mean and SD.

### Exploring other ways to promote artificial chromoplast differentiation

We next explored the possibility of increasing leaf β-carotene contents by promoting crtB-mediated artificial chromoplastogenesis following a pharmacological and a genetic strategy. For the former, we reasoned that the exogenous application of phytohormones could positively (or negatively) impact the formation of artificial chromoplasts resulting in higher (or lower) β-carotene levels. We applied the phytohormones with a fine brush on the surface of *N. benthamiana* leaves 2 h after agroinfiltration with the p-crtB construct (i.e., once the agroinfiltration halo was dried). Then, ɸPSII was measured every 12 h from 24 to 72 hpi as a reliable physiological marker to monitor the speed of chloroplast-to-chromoplast differentiation, as described above (Llorente et al., 2020), and carotenoid levels were quantified at 96 hpi (Figure 3). Most plant hormones had no effect on ɸPSII drop rate compared to leaves treated with a mock solution (Figure 3A). The only exceptions were gibberellins (gibberellic acid, GA_3_), which accelerated the chromoplastogenesis process, and synthetic strigolactones (GR24) and auxins (picloram), which delayed it (Figure 3A). In particular, the application of GA_3_ resulted in a significant decrease of ɸPSII in p-crtB samples at 36 hpi compared to the mock control (Figure 3A), eventually leading to a higher content of carotenoids, including β-carotene (Figure 3B). By contrast, leaves treated with GR24 or picloram showed significantly higher φPSII from 48 dpi (Figure 3A) and lower levels of carotenoids at 96 hpi (Figure 3B). To test whether the internal amounts of these phytohormones could regulate the chromoplastogenesis process, we used inhibitors of the endogenous biosynthesis pathways. Specifically, we used paclobutrazol to block endogenous gibberellin biosynthesis (Heden and Graebe, 1985), the 4-hydroxyaryl hydroxamic acid D2 to inhibit strigolactone biosynthesis (Harrison et al., 2015), and L-kynurenine to inhibit auxin biosynthesis (He et al., 2011). However, none of these inhibitors had any significant effect on φPSII or carotenoid contents compared to plants treated with the mock solution (Figure 3). While the GA_3_ treatment did slightly improve strategy P, it had deleterious effects in non-agroinfiltrated leaves and stems, which turned longer and paler. Also, the phytohormone treatment had to be performed in a specific moment after agroinfiltration (to avoid the formation of too much humidity on the agroinfiltration halo that could affect the efficiency of transformation), which makes it difficult to implement in large-scale production settings. These factors, together with regulatory constraints associated with the use of chemicals and the high economic cost of the treatment, discarded treating with gibberellins as a β-carotene biofortification method for commercial use.

**Figure 3.**
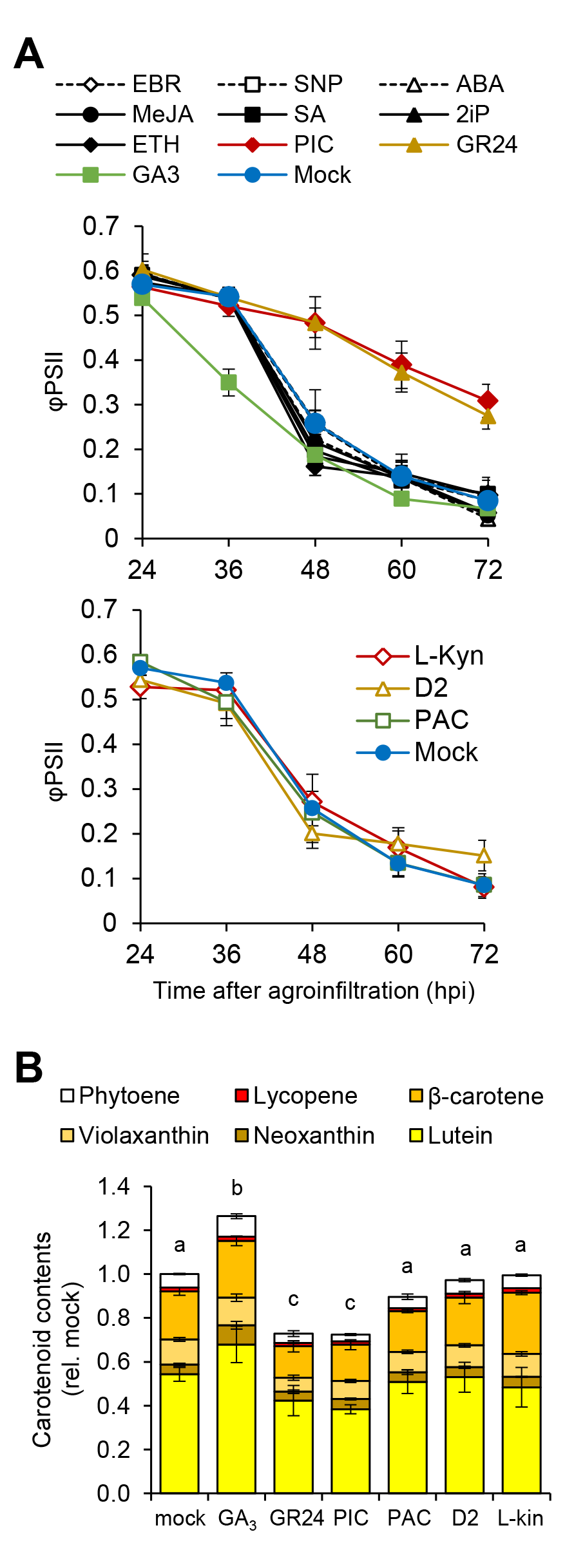
Treatment with some phytohormones interferes with the chromoplastogenesis process. *N. benthamiana* leaves agroinfiltrated with the p-crtB construct were treated with the indicated compounds or a mock solution. **(A)** Effective quantum yield of photosystem II (φPSII) values in leaf areas treated with the indicated phytohormones (upper plot), or inhibitors (lower plot). Hormone treatments that had no effect compared to the mock treatment are shown in black in the upper plot. Mean and SD of at least 5 measurements in n≥3 independent leaf samples are shown. **(B)** Relative carotenoid levels at 96 hpi shown as the mean and SD of n≥3 independent biological replicates. Letters represent statistically significant differences (*P* < 0.05) among means of total carotenoid levels according to post hoc Tukey’s tests run after one way ANOVA detected different means. See Table 1 for acronyms.

**Table 1.** Compounds applied to test hormonal regulation of artificial chromoplast development in *N. benthamiana* leaves. All the compounds were diluted in 0.05 % Tween20 (in water) to the indicated concentration before application to the leaf surface with a fine brush.

| Class | Type | Compound | Acronym | Concentration ($\mu\text{M}$) |
| --- | --- | --- | --- | --- |
| Hormone | Brassinosteroid | Epibrassinolide | EBR | 10 |
|  | Cytokinin | 2-isopentenyladenine | 2iP | 10 |
|  | Absciscic acid | Absciscic acid | ABA | 10 |
|  | Jasmonate | Methyl jasmonate | MeJA | 100 |
|  | Salicylic acid | Salicylic acid | SA | 300 |
|  | Gibberellin | Giberellic acid | GA3 | 100 |
| Synthetic hormone | Strigolactone analog | GR24 | GR24 | 100 |
|  | Auxin analog | Picloram | PIC | 50 |
| Hormone donor | Nitric oxide donor | Sodium nitroprusside | SNP | 100 |
|  | Ethylene donor | Etephon | ETH | 1500 |
| Hormone synthesis inhibitor | Gibberellin inhibitor | Paclobutrazol | PAC | 10 |
|  | Strigolactone inhibitor | 4-hydroxyaryl hydroxamic acid | D2 | 100 |
|  | Auxin inhibitor | L-Kynurenine | L-Kyn | 500 |

We next carried out a genetic screen of Arabidopsis mutants with an altered plastid-to-nucleus (i.e., retrograde) communication, such as *ron1*/*sal1* (Estavillo et al., 2011), *csb3/ceh1* (Xiao et al., 2012) and *gun1* (Koussevitzky et al., 2007). The rationale was that defects in particular retrograde signaling pathways might interfere with chromoplastogenesis, hence providing hints on molecular mechanisms that could eventually be manipulated to optimize the differentiation process. Chromoplastogenesis was triggered in Arabidopsis wild-type and mutant plants using viral vectors to deliver crtB to leaf cells (Llorente et al., 2020; Morelli et al., 2023). Soil-grown plants were infected with the viral vector TuMV-crtB and the levels of carotenoids were measured 3 weeks later, when symptoms (leaves turning vivid yellow) were obvious. None of the mutants analyzed showed significant changes in their carotenoid profile or β-carotene content compared to wild-type controls (Figure 4), indicating that chloroplast-to-chromoplast differentiation in Arabidopsis leaves can proceed normally despite the defective operation of at least some retrograde signaling pathways.

**Figure 4.**
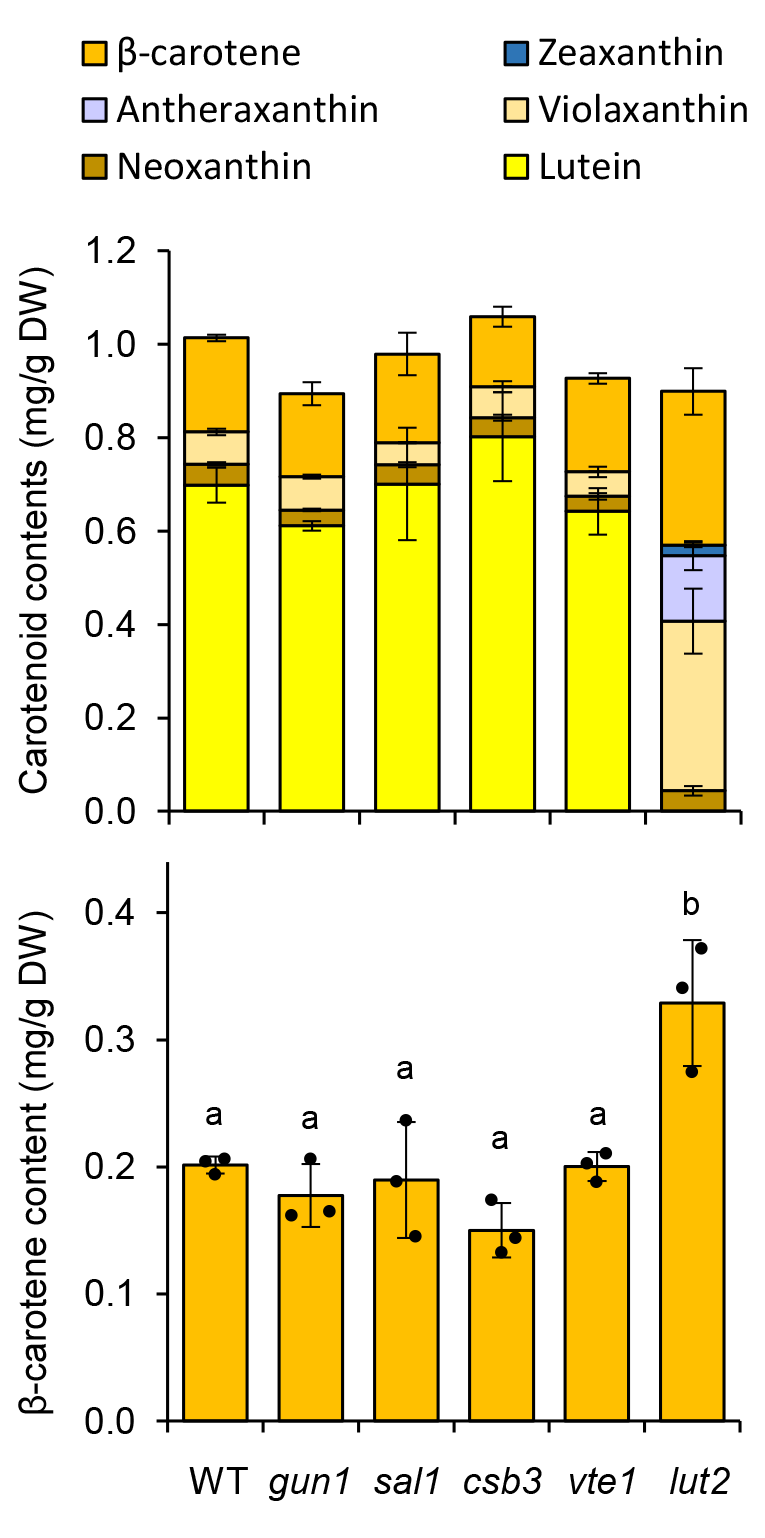
Genetic manipulations of retrograde and carotenoid-related pathways hardly impact β- carotene levels. Arabidopsis wild-type (WT) plants and the indicated mutants were infected with the TuMV-crtB vector. After 3 weeks, rosette leaves showing the phenotypes associated with viral infection (e.g., curled leaves) and ctrB activity (e.g., yellow leaves) were collected to measure carotenoid contents. Plots represent mean and SD of total carotenoid levels (upper) and β-carotene contents (lower) of n=3 samples from different plants. Dots represent individual values and letters represent statistically significant differences (*P* < 0.05) among means according to post hoc Tukey’s tests run after one way ANOVA detected different means.

Finally, we tested a genetic strategy to increase β-carotene content based on the use of mutants in competing biosynthetic steps. In particular, we used the tocopherol biosynthesis mutant *vte1* (Sattler et al., 2003) to decrease competition for GGPP and the lycopene ε cyclization mutant *lut2* (Pogson et al., 1996) to block the β,ε branch of the carotenoid pathway leading to lutein and hence divert all precursors to the β-carotene branch (Figure 1A). Upon infection with the TuMV-crtB vector, wild-type and mutant plants showed no significant differences in total carotenoid levels in yellow (i.e., chromoplast-containing) leaf tissues (Figure 4). In the case of the *lut2* mutant, however, we did detect a modest (1.5-fold) enrichment in β-carotene contents (Figure 4). Higher enrichments would be expected by blocking the conversion of β-carotene into downstream xanthophylls (Figure 1A), which accumulate at high levels in the chromoplasts of *lut2* leaves (Figure 4).

### High-light exposure is an effective strategy to further enrich leaves in β-carotene

Artificial chromoplasts in crtB-producing leaves accumulate phytoene and β-carotene in PG, where most of the crtB protein is localized (Morelli et al., 2023). Additionally, PG from artificial chromoplasts harbor higher levels of isoprenoid vitamins E (tocopherols) and K1 (phylloquinone), making them interesting structures to target for nutritional enrichment of leaves (Morelli et al., 2023). Among the environmental cues that result in a high proliferation of PG, high-light exposure is particularly interesting because, besides increasing PG amount and size, it also affects stimulates the accumulation of pigments with photoprotective function (Lichtenthaler, 2007). We therefore tested the effect of increased light intensity on the accumulation of β-carotene in HEBIY samples (Figure 5). *N. benthamiana* plants grown under normal light conditions (50 μmol photons·m^−2^·s^−1^ white light, namely W50) were exposed for 3 days to 10-fold higher light intensity (500 μmol photons·m^−2^·s^−1^ white light, W500) or kept under W50. Then, leaves from the two sets of plants were agroinfiltrated with the HEBIY combination, and plants were kept under W50 until samples were taken for HPLC analysis at 6 dpi (Figure 5). As expected (Lichtentaler et al., 2007), leaves pre-treated with high-light (W500) showed a higher content of total carotenoids compared to leaves sampled from control plants grown under lower irradiances (W50). In particular, increased levels of β-carotene were observed while other carotenoids (e.g., violaxanthin) were reduced (Figure 5A), similar to our previous results with p-crtB leaves (Morelli et al., 2023). As a summary, combining strategies C and P with high-light treatment resulted in almost a 3-fold increase in total carotenoid levels but a more than 7-fold increase in β-carotene (pro-vitamin A) contents of *N. benthamiana* leaves compared to controls agroinfiltrated with the GFP protein (Figure 5B). Indeed, the proportion of β-carotene relative to total leaf carotenoids was raised from around 20% in the chloroplasts of GFP control leaves to 30% in the chromoplasts of crtB samples and 45% in HEBIY leaves exposed to high-light (Figure 5B).

**Figure 5.**
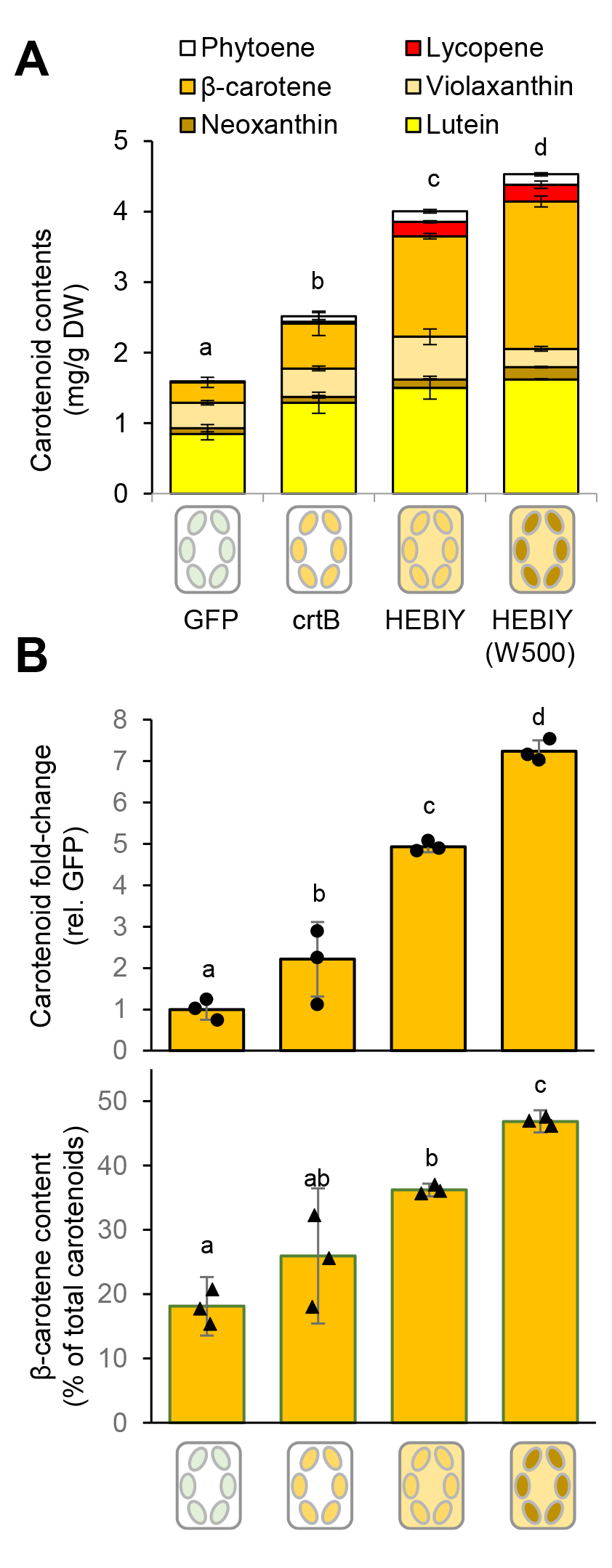
Exposure to high irradiances further increases β-carotene levels. *N. benthamiana* plants grown under normal light conditions (W50) were exposed for 3 days to 10-fold higher light intensity (W500) or kept under W50. Then, leaves from the two sets of plants were agroinfiltrated with the indicated constructs and analyzed at 6 dpi. **(A)** Absolute carotenoid levels shown as the mean and SD of n=3 independent biological replicates. **(B)** Relative levels of β-carotene in the samples shown in (A). The upper plot represents the contents relative to those in control (GFP) samples, whereas the lower plot shows the proportion relative to total carotenoids in every combination. Dots represent individual values and letters indicate statistically significant differences (*P* < 0.05) among means according to post hoc Tukey’s tests run after one way ANOVA detected different means.

### Accumulation of β-carotene in PG and extraplastidial locations increases its bioaccessibility

Despite the remarkable increase in both the levels and the proportion of β-carotene in leaves achieved by our strategies, enriching carotenoid contents in a particular food is only the first step towards efficient biofortification (Morelli and Rodriguez-Concepcion, 2023). Carotenoids are lipophilic isoprenoids that must first be released from the food matrix and then incorporated into water-miscible intestinal micelles. The fraction of a nutrient (e.g., a carotenoid) accessible for absorption in this condition is referred to as bioaccessibility. The physicochemical contexts and subcellular locations where carotenoids accumulate in plant cells highly influence bioaccessibility (Watkins and Pogson, 2020; Zheng et al., 2020). In the case of β-carotene, its bioaccessibility in lettuce artificial chromoplasts produced by crtB was doubled compared to that in chloroplasts (Morelli and Rodriguez-Concepcion, 2022). Recent results showed that extraplastidial β-carotene produced with a fungal pathway engineered in plant cells accumulates in cytosolic lipid droplets (Zheng et al., 2023), an environment that might be beneficial for bioaccessibility. To test whether the extraplastidial β-carotene produced with our bacterial pathway might be more bioaccessible, we performed an *in vitro* assay developed in our lab (Morelli and Rodriguez-Concepcion, 2022) to estimate the bioaccessibility of carotenoids in leaf samples. *N. benthamiana* leaves were agroinfiltrated with HE (a control for β-carotene exclusively present in chloroplasts), p-crtB (in which β-carotene is stored in the PG of artificial chromoplasts), HEcBIY (with β-carotene accumulated in chloroplasts but also in extraplastidial locations) or HEBIY (in which β-carotene is accumulated in both artificial chromoplasts and extraplastidial locations). At 6 dpi, samples were collected and used to estimate the percentage of carotenoids remaining in the tissue after *in vitro* digestion (Figure 6). We observed that bioaccessibility of the β-carotene engineered in the cytosol (HEcBIY) was only marginally higher compared to that produced and stored in chloroplasts (HE). Interestingly, induction of chromoplastogenesis by plastidial crtB significantly increased bioaccessibility of β-carotene (Figure 6). Because β-carotene in chloroplasts locates in membranes whereas it is mostly associated with PG in artificial chromoplasts (Morelli et al., 2023), we reasoned that its accumulation of these plastidial lipid bodies might explain the increase in bioaccessibility. To test this hypothesis, we promoted PG proliferation by exposure to high-light (W500) similarly to that described above and then calculated bioaccessibility in p-crtB and HEBIY leaves at 6 dpi (Figure 6). W500 treatment resulted in increased β-carotene bioaccessibility in both p-crtB and HEBIY samples, but it was statistically significant only in the case of p-crtB, likely because the bioaccessibility of the large amount of extraplastidial β-carotene found in HEBIY samples hardly changes (Figure 6). The conclusion that PG localization enhances β-carotene bioaccessibility is indirectly supported by the unchanged bioaccessibility of lutein in PG-enriched W500 samples compared to W50 controls, since lutein does not accumulate in the PG found in artificial leaf chromoplasts (Morelli et al., 2023). Considering both the amount and the bioaccessibility, the combination of strategies C and P together with the high-light treatment (HEBIY + W500) resulted in an impressive 30-fold increase in the contents of accessible β- carotene compared to HE controls (Figure 6).

**Figure 6.**
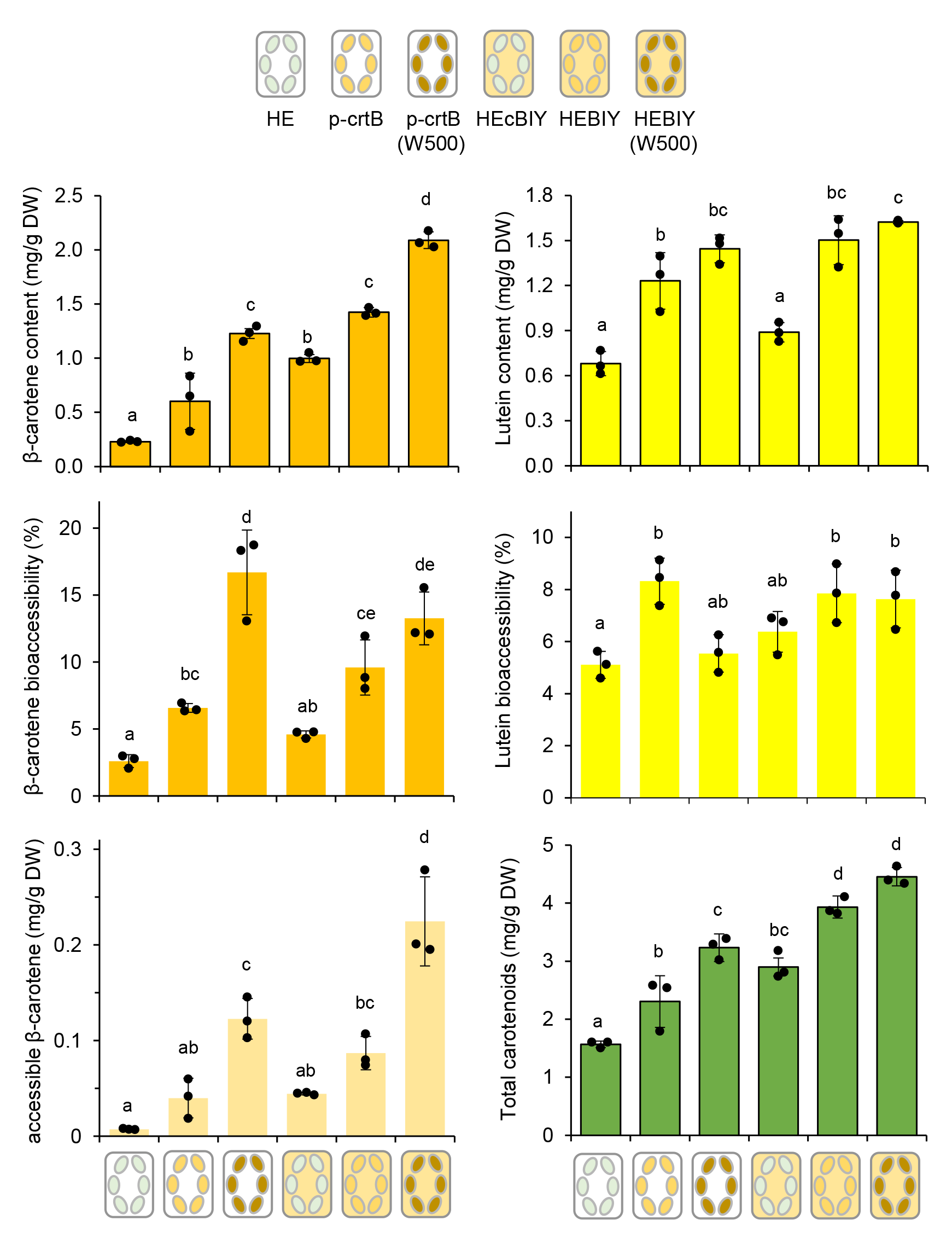
Subcellular accumulation impacts β-carotene bioaccessibility. *N. benthamiana* plants grown under normal light conditions (W50) were exposed for 3 days to 10-fold higher light intensity (W500) or kept under W50. Then, leaves from the two sets of plants were agroinfiltrated with the indicated constructs and analyzed at 6 dpi. Plots show carotenoid levels, bioaccessibility, and accessible content as the mean and SD of n=3 independent biological replicates. Dots represent individual values and letters indicate statistically significant differences (*P* < 0.05) among means according to post hoc Tukey’s tests run after one way ANOVA detected different means.

### Developed strategies can be applied to the biofortification of edible lettuce leaves

The reported results illustrate how triggering artificial chromoplastogenesis in *N. benthamiana* leaves either with crtB alone (strategy P) or in combination with a cytosolic pathway (strategy C) and/or exposure to high-light can substantially increase not only the contents but also the bioaccessibility of β- carotene, the main pro-vitamin A carotenoid. Our previous work demonstrated that strategy C could be implemented in lettuce, a plant producing edible green leaves (Andersen et al. 2021). We also showed that delivery of crtB to leaf cells using viral vectors is able to promote the conversion of chloroplasts into chromoplasts in all plants tested (Majer et al., 2017; Llorente et al., 2020; Houhou et al., 2022; Rodriguez-Concepcion and Daròs, 2022). We therefore investigated whether the described nutritional benefits of developing artificial chromoplasts and promoting PG proliferation could also be extended to lettuce. The viral vector LMV-crtB or an empty LMV control (Llorente et al., 2020) were agroinfiltrated in leaves of the Romaine variety of lettuce (Figure 7). About 2 weeks later, LMV-crtB leaves had developed a characteristic orange/yellow color and a strong increase in their carotenoid content compared to control LMV leaves, which showed a normal green color but very curly leaves as expected from viral infection (Figure 7A). The characteristic lettuce carotenoid lactucaxanthin also increased in LMV-crtB samples, confirming a general upregulation of the whole carotenoid pathway (Figure 7B). Similar to that described in the *N. benthamiana* system (Morelli et al., 2023), crtB also promoted the development of PG in lettuce, as deduced from the increased levels of PG-associated fibrillins FBN1a and FBN2 in LMV-crtB samples compared to LMV controls (Figure 7C).

**Figure 7.**
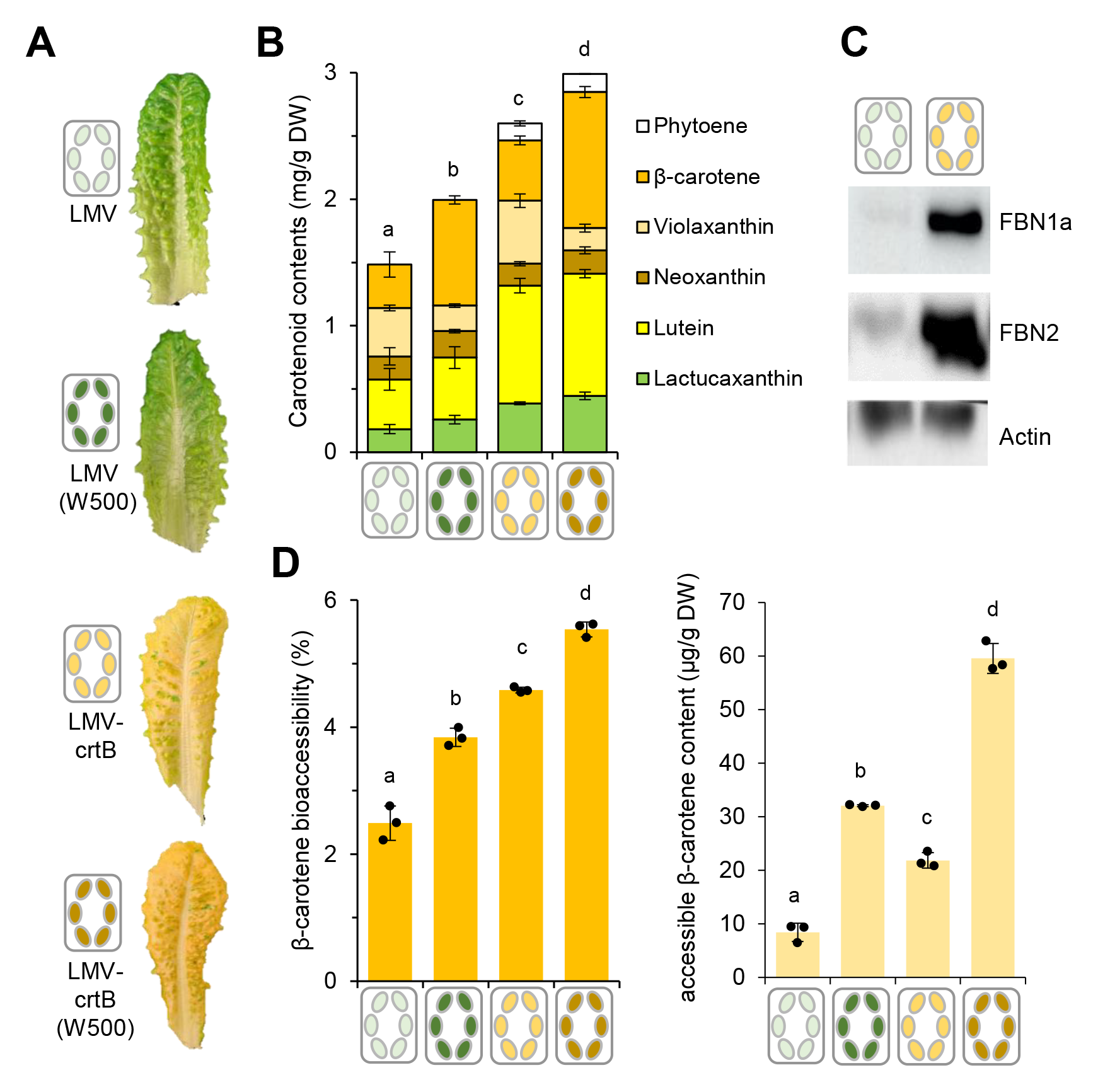
High-light treatment increases β-carotene contents and bioaccessibility in lettuce leaves. Romaine lettuce plants grown under normal light conditions (W50) and inoculated with the indicated viral vectors were exposed for 5 days to 10-fold higher light intensity (W500) or kept under W50. After transferring them back to W50 for 2 days for recovery, leaves from the two sets of plants were collected and snap-frozen for further analysis. **(A)** Representative images of collected leaves with a schematic presentation of treatments. **(B)** Plot representing the mean and SD of carotenoid levels in n=3 independent biological replicates per treatment. **(C)** Immunoblot analysis of PG-associated protein markers (fibrillins FBN1a and FBN2) in the indicated samples. Actin levels were analyzed as loading controls. **(D)** Bioaccessibility and accessible content of β-carotene in the samples analyzed in (B). Dots represent individual values and letters indicate statistically significant differences (*P* < 0.05) among means according to post hoc Tukey’s tests run after one way ANOVA detected different means.

We next-tested the effect of high-light exposure on carotenoid accumulation in lettuce. In this case, infection with the viral vectors (either LMV or LMV-crtB) was carried out first and, at 6 dpi (i.e., when the first yellow spots started to develop on the leaves), plants were exposed to W500 or left under W50 as a control for 5 more days. After a 2-day recovery period under W50, leaf samples were collected and used to estimate carotenoid contents and β-carotene bioaccessibility (Figure 7D). Again coherently with what was observed in *N. benthamiana* leaves (Figure 6), high-light caused an increase in total carotenoids in general and β-carotene in particular (Figure 7B). Interestingly, such carotenoid biofortification also took place in control leaves not expressing a foreign protein such as crtB (i.e., harboring chloroplasts). Furthermore, β-carotene bioaccessibility also improved substantially in leaves exposed to high-light regardless of whether they were harboring chloroplasts (LMV) or chromoplasts (LMV-crtB), even though the highest levels were detected when crtB and high-light treatments were combined (Figure 7B). In terms of increasing accessible β-carotene contents, the treatment with high-light worked better than the use of crtB (strategy P), and the highest levels were achieved when combining both approaches (Figure 7D).

## DISCUSSION

Biofortification of green leafy vegetables with pro-vitamin A carotenoids is still in its infancy, in part because of the challenges associated with changing the balance between carotenoids and chlorophylls in chloroplasts and hence negatively impacting photosynthesis (Domonkos et al., 2013; Esteban et al., 2015; Hashimoto et al., 2018; Zhen and van Iersel, 2017; Zheng et al., 2020). Here we present two strategies that can be used separately or combined to overcome this problem. Strategy C involves producing pro-vitamin A carotenoids such as β-carotene in extraplastidial locations to avoid disrupting the balance of photosynthetic pigments and hence interfering with photosynthesis. Strategy P was implemented by triggering the conversion of chloroplasts into chromoplasts only in some leaves at the end of their development, leaving other leaves with active chloroplasts to sustain normal photosynthesis and plant functions. We further demonstrate that promoting PG proliferation with strategy P but also with high-light treatment increases β-carotene bioaccessibility in edible lettuce leaves, unveiling the power of non-GMO approaches for leaf biofortification.

The strategies presented here are just a proof of concept, and they will require future improvements to exploit their full potential. In the case of strategy C, i.e., the possibility to produce carotenoids in compartments other than plastids, the 2-fold increase in total carotenoid levels and 4-fold increase in β-carotene contents confirms a remarkable capacity of leaf cells to accumulate carotenoids outside chloroplasts. The precise location where extraplastidial β-carotene is synthesized and the form in which it is stored in the cytosol is unknown, but recent results suggest that pro-vitamin A carotenoids produced in the cytosol of *N. benthamiana* leaf cells accumulate in lipid droplets (Zheng et al., 2023). In any case, our data suggest that factors besides storage might be more limiting to further increase the contents of this pro-vitamin A carotenoid. In particular, high levels of residual lycopene and phytoene that are not converted into β-carotene (Figure 1B) suggest that crtI and crtY activities are important limiting steps. These bacterial enzymes likely interact to form a multiprotein complex that facilitates substrate channeling (Nogueira et al., 2013; Ravanello et al., 2003). Engineering a scaffolding system to get all the carotenoid mini-pathway enzymes together might contribute to improve substrate delivery and product release, eventually resulting in higher β-carotene titers. Alternatively, the use of directed evolution to create crtI and crtY enzymes optimized to work in a plant cytosolic environment might contribute to improve the corresponding enzymatic conversions.

A bottleneck has also been unveiled in the case of strategy P. Upregulation of the MEP pathway flux by overexpressing the tomato *SlDXS1* gene resulted in a high increase in the level of phytoene but no significant increase in downstream carotenoids (Figure 2). Besides confirming that SlDXS1 works to increase the MEP pathway flux, this result also suggested that the endogenous GGPP synthase enzymes could efficiently convert the extra supply of MEP-derived IPP and DMAPP into GGPP and that the plastid-localized crtB enzyme was able to convert enhanced levels of GGPP into phytoene. However, the conversion of the extra phytoene into downstream carotenoids appeared to be limiting in *N. benthamiana* leaves. Several explanations for the observed poor phytoene conversion are possible. It is likely that biosynthetic and storage mechanisms could already be saturated in crtB-containing samples, thus limiting further increases upon enhancing the supply of phytoene. Most crtB-derived phytoene is produced and stored in PG, where it can be further converted into β-carotene (Morelli et al., 2023). However, it is also possible that the additional phytoene produced when SlDXS1 is added cannot be accessed by endogenous desaturase enzymes due to differential compartmentation. It is intriguing that only the tomato SlDXS1 isoform worked to produce more phytoene. In the case of the AtDXS enzyme, it can be argued that the SlDXS1 enzyme comes from tomato, and hence might work better in another species of the Solanaceae family - such as *N. benthamiana* - than an enzyme from the Brassicaceae family. In agreement with this possibility, the production of phytoene in Golden rice (monocot) was highly increased when a monocot PSY (from maize) was used instead of the initial dicot PSY (from daffodil) enzyme (Paine et al., 2005). Alternatively, AtDXS and/or SlDXS2 might have a lower catalytic activity than SlDXS1, or they might interact with different partners. For example, it has been shown that the tomato chromoplast-associated PSY1 isoform is much more active than the chloroplast-associated PSY2 (Cao et al., 2019), and these two enzymes can directly interact with the GGPP synthase isoform SlG2 but not with SlG3 (Barja et al., 2021). It is therefore possible that the presence of SlDXS1 somehow results in the production of phytoene in a compartment different from that where desaturases and isomerases can transform it into lycopene. In any case, the upregulated production of phytoene was found to speed up the φPSII drop associated with the initial loss of chloroplast identity (Figure 2B), consistent with the proposed model that the accumulation of this carotenoid precursor is the main trigger of the artificial chromoplast differentiation process (Llorente et al., 2020).

The additive phenotype resulting from combining strategies C and P (Figure 1B) demonstrates that the cytosolic MVA pathway and the plastidial MEP pathway can work together at the same time without downregulating (or upregulating) each other. This information should open the door to engineer the production of isoprenoids of interest in plant biofactories by using not one but the two pathways supplying their metabolic precursors. The conversion of leaf chloroplasts into chromoplasts in strategy P, however, involves a loss of photosynthetic activity that restricts its application. The use of agroinfiltration, viral vectors, or specific promoters allows to target specific leaf areas, tissues or developmental stages while preserving the photosynthetic capacity of the rest to support photosynthesis. Both agroinfiltration and viral vectors have been previously demonstrated to work in lettuce to produce extraplastidial carotenoids by strategy C (Andersen et al., 2021) and to promote carotenoid accumulation by differentiating artificial chromoplasts by strategy P (Llorente et al., 2020; Morelli and Rodriguez-Concepcion, 2022). Here we go a step forward by showing that both the accumulation and the bioaccessibility of β-carotene increase after inducing the proliferation of PG in lettuce leaves either by strategy P or by just exposing unmodified leaves to high-light (Figure 7). Lettuce is an excellent source of vitamins, minerals, and bioactive compounds such as polyphenols and carotenoids (Shi et al., 2022). Although a strong influence of light and other environmental conditions on the contents of these health-promoting phytonutrients has been reported, the positive effect of high-light and PG differentiation on β-carotene accumulation and bioaccessibility was unknown. A correlation between PG storage and bioaccesibility has been shown for β-carotene and other carotenoids in several plant systems. For example, fruit containing abundant PG such as mango and papaya showed a higher β-carotene bioaccesibility than fruits with chromoplasts with few or no PG such as melon or grapefruit (Jeffery et al., 2012a, 2012b). In the chloroplasts of green tissues, β- carotene is associated with the photosynthetic apparatus in tightly built pigment-protein complexes resulting in a more difficult release during digestion and a subsequent lower bioaccessibility compared to the pigment found in chromoplasts (Jeffery et al., 2012a, 2012b; Lichtenthaler, 2007). Promoting PG proliferation with a high-light treatment (Van Wijk and Kessler, 2017) likely facilitates the accumulation of β-carotene in these subcompartments and hence improves bioaccessibility.

Our data demonstrate that impressive (about 30-fold) increases in accessible β-carotene contents are feasible in leaves (Figure 6) and that the generated knowledge can be steadily translated to biofortify green leafy vegetables such as lettuce (Figure 7). Improving the levels and storage of carotenoids in lettuce has the added advantage of providing an increased amount of lactucaxanthin, a unique ε,ε- carotenoid reported to have antidiabetic and antioxidant properties (Gopal et al., 2017). However, our pharmacological (Figure 3) and genetic (Figure 4) approaches to learn about the molecular mechanisms supporting artificial chromoplastogenesis did not provide meaningful insights. Further efforts (i.e., dynamic transcriptomic and proteomic analyses) would be required to generate such knowledge and then apply it to improve strategy P. Furthermore, there is still a long way to the commercial application of this technology. While advances in synthetic biology are permitting to deploy complex gene circuits in plants that activate upon chemical or physical treatments at appropriate tissues or developmental stages, reaching the market will strongly depend on the evolution of societal and political concerns towards genetic manipulation (Rodriguez-Concepcion and Daros, 2021). Future developments to deliver nucleic acid sequences and proteins into plant cells without using bacterial or viral vectors may solve at least some of these limitations.

### EXPERIMENTAL PROCEDURES

#### Plant material and growth conditions

*Nicotiana benthamiana* and *Lactuca sativa* (var. Romaine) plants used for the transient expression assays were grown in a greenhouse under a stable photoperiod of 14 h of light (50 µmol m^-2^ s^-1^ photons, referred to as W50) at 26°C and 10 h dark at 21°C. For high-light treatment, 4-week-old *N. benthamiana* plants were moved to an Aralab Fitoclima 600 plant growth chamber with the same photoperiod but higher light intensity (500 µmol m^-2^ s^-1^ photons, W500) for 3 days. After agroinfiltration plants were maintained at W50. In the case of lettuce, 4-week-old plants were inoculated with *N. benthamiana* tissue infected with a viral vector and 6 days later they were moved to W500 conditions for 5 days. Then, plants were moved to W50 conditions for 2 additional days for recovery. For treatments with phytohormones and inhibitors, we diluted them in water and 0.05 % Tween 20 (to lower surface tension and delay the evaporation of the solution) at the concentrations indicated in Table 1.

#### Gene constructs and transient expression assays

Constructs overexpressing tHMGR, crtE, crtI and the different versions of crtB used in this study for the assembling of the extraplastidial pathway were available in the lab (Llorente et al., 2020; Andersen et al. 2021). The *P. ananatis crtY* gene was cloned in the Gateway pGWB405 vector as described (Llorente et al., 2020). Similarly, cDNA sequences for AtDXS, SlDXS1, and SlDXS2 were amplified from Arabidopsis or tomato and cloned into pGWB405 (AtDXS), pGWB420 (SlDXS1) or pGWB454 (SlDXS2).

Agroinfiltration assays were carried out as described (Andersen et al. 2021) using *Agrobacterium tumefaciens* GV3101 strains transformed with constructs of interest. Cultures were mixed in identical proportions for the various combinations. For the inoculation of lettuce, 3 leaves of 3-4 weeks old plants were infiltrated in at least 6 different spots each with the same solution used for *N. benthamiana*, without the use of a gene silencing helper. Inoculation of Arabidopsis plants with viral vectors was performed as described (Llorente et al., 2020).

#### Photosynthetic measurements

Effective quantum yield of PSII (ɸPSII) was measured with a MAXI-PAM fluorimeter (Heinz Walz GmbH) or a Handy GFP Cam (PSI (Photon Systems Instruments) as (Fm’-Fs)/Fm’, where Fm’ and Fs are the maximum and minimum fluorescence of light-exposed plants, respectively. The chosen light intensity was 21 PAR (AL=2). Average values were calculated from three biological replicates with three different leaf areas for each replicate.

#### In vitro bioaccessibility assay

The *in vitro* digestion procedure was performed as described (Morelli and Rodriguez-Concepcion, 2022). The digested and undigested samples were analyzed by HPLC as described above. Bioaccessibility was calculated as the ratio between carotenoid concentration after digestion and the concentration of the same compound in the starting undigested sample. Accessible carotenoid contents were calculated by multiplying the amount of carotenoid in the sample by the corresponding bioaccessibility percentage.

#### Protein and metabolite analysis

Carotenoids and chlorophylls were extracted and analyzed by HPLC-DAD as described (Barja et al., 2021). Protein extraction and separation were carried out as described (Morelli et al., 2013) using approximately 5 mg of freeze-dried leaf tissue. For immunoblot analysis, primary antibodies against FBN1A, FBN2, and actin and a secondary antibody conjugated with horseradish peroxidase (Millipore) were used. Detection of immunoreactive bands was performed using SuperSignal West Pico PLUS (Thermo Scientific). Chemiluminescent signals were visualized using an ImageQuant 800 biomolecular imager (Amersham).

## ACKNOWLEDGEMENTS

We thank Trine B. Andersen and Jose A. Daros for materials and M. Rosa Rodriguez, Jose Perez-Beser and the staff at the IBMCP Metabolomics Platform for technical support. This work was funded by grants from Spanish MCIN/AEI/10.13039/501100011033 and European NextGeneration EU/PRTR and PRIMA programs to MR-C (PID2020-115810GB-I00 and UToPIQ-PCI2021-121941). MR-C is also supported by Generalitat Valenciana (PROMETEU/2021/056 and AGROALNEXT/2022/067) and the MCIN/AEI-funded Spanish Carotenoid Network, CaRed (RED2022-134577-T). LM and PP-C received predoctoral fellowships from La Caixa Foundation (INPhINIT program LCF/BQ/IN18/11660004) and Generalitat Valenciana (CIACIF/2021/278), respectively. We declare no conflict of interest.

